# Genome-wide profiling of heritable and *de novo* STR variations

**DOI:** 10.1101/077727

**Authors:** Thomas Willems, Dina Zielinski, Assaf Gordon, Melissa Gymrek, Yaniv Erlich

**Affiliations:** New York Genome Center, New York, NY 10013, USA; Computational and Systems Biology Program, MIT, Cambridge, MA 02139, USA; Analytic and Translational Genetics Unit, Massachusetts General Hospital, Boston, MA, USA; Program in Medical and Population Genetics, Broad Institute of MIT and Harvard, Cambridge, MA, USA; Department of Computer Science, Fu Foundation School of Engineering, Columbia University, New York, NY 10027, USA; Center for Computational Biology and Bioinformatics, Columbia University, New York, New York 10032, USA

## Abstract

Short tandem repeats (STRs) are highly variable elements that play a pivotal role in multiple genetic diseases, population genetics applications, and forensic casework. However, STRs have proven problematic to genotype from high-throughput sequencing data. Here, we describe HipSTR, a novel haplotype-based method for robustly genotyping, haplotyping, and phasing STRs from whole genome sequencing data and report a genome-wide analysis and validation of *de novo* STR mutations.

---

The impact of genomics is contingent upon its ability to identify genetic variants. While tremendous progress has been made in identifying nearly every type of genetic variation, short tandem repeat (STR) variations remain largely understudied. Composed of repeating 1-6 base pair motifs, STRs are among the most polymorphic variants in the human genome and are present at over 1 million loci. STRs play a key etiological role in more than 30 Mendelian disorders^1^ and recent evidence has underscored their profound regulatory role and potential involvement in complex traits^2–4^. Beyond medical genetics, STRs have proven useful for various applications in population genetics, forensics, and single cell lineage analysis. Despite their utility, STRs are typically omitted from large-scale genetic studies. While several STR callers exist^5–7^, they suffer from limited accuracy, difficulties in calling homopolymer runs, sensitivity to STR stutter noise, and limited functionality compared to mature SNP callers. As a result, large-scale projects^8–10^ are reluctant to report STR genotypes and are essentially blind to many of the most variable parts of the genome.

Here, we developed a novel algorithm called HipSTR (haplotype-inference and phasing for STRs) to create a mature tool for STR studies. HipSTR builds on our extensive experience with STR genotyping and addresses major limitations of existing STR tools^5,11,12^. To improve accuracy, our tool uses a multitude of inference techniques that integrate additional information about the haplotype in which the STR resides (Supplementary Fig. 1). Briefly, HipSTR begins by learning a parametric model that captures the stutter noise profile specific to each locus. Using prior information about the genomic location of the repeat, it then harnesses this model to tune a hidden Markov model (HMM) that carefully realigns the STR-containing reads and mitigates the effects of stutter and sequencing errors (Supplementary Figs. 1-2). The realignment framework is highly flexible and can integrate population-scale data from other individuals and phased SNP scaffolds to find the most likely set of alleles, conferring robustness to the genotyping process (Supplementary Figs. 3-4). The output of HipSTR is a VCF file that consolidates all variants at an STR locus into a single line. The tool is written in C++ and is freely available at https://hipstr-tool.github.io/HipSTR.

**Figure 1:**
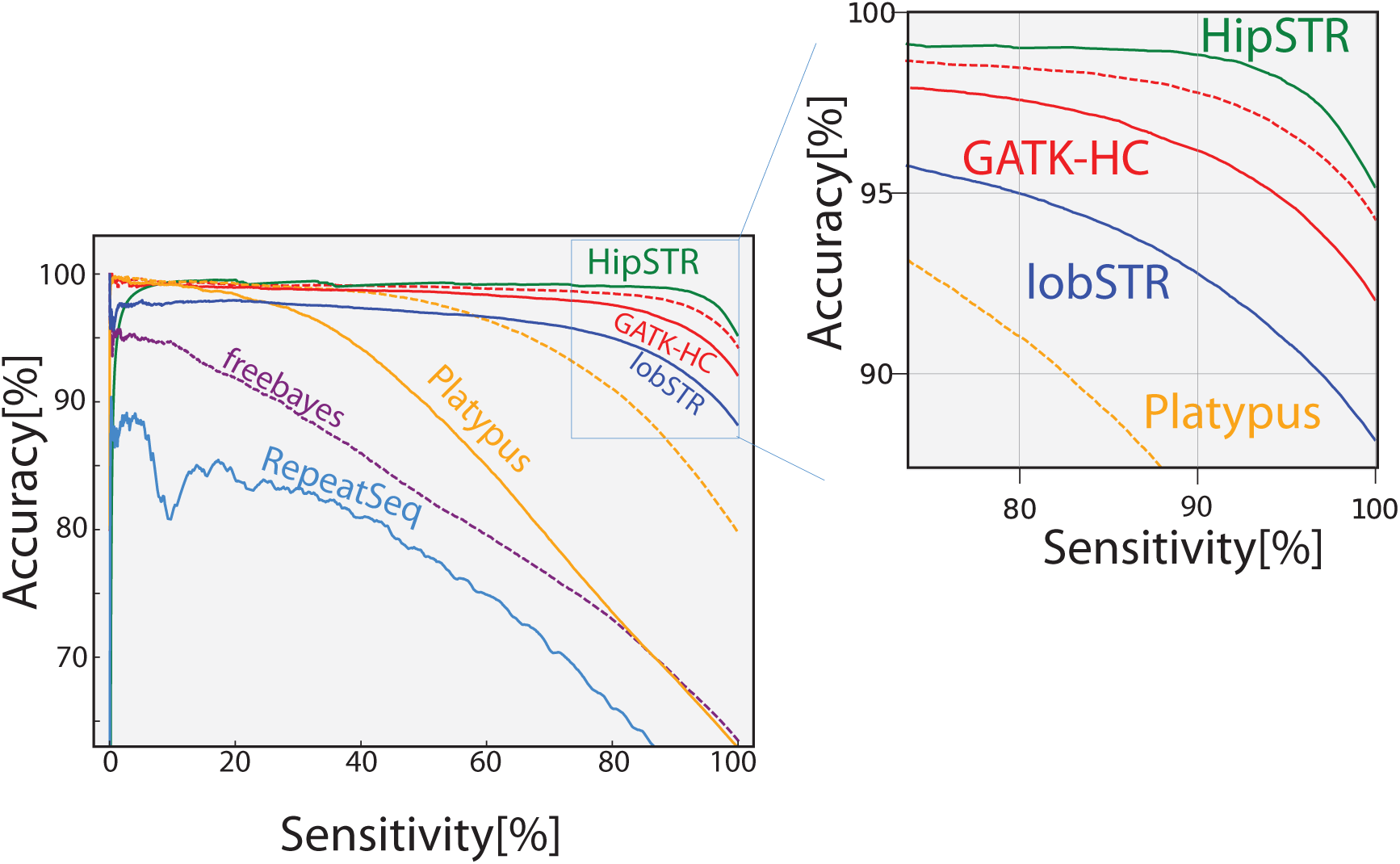
STR genotype accuracy. We tested the accuracy of each variant caller to genotype STRs from the Marshfield panel. After generating calls for each tool, we ranked the confidence in the STR genotypes using a machine-learning classifier and explored the accuracy as a function of sensitivity (number of STR variants that crossed a certain confidence threshold). In all relevant circumstances, HipSTR’s classifier resulted in the most accurate STR genotypes. Solid lines denote tools run using default settings, while dashed lines denote tools run using settings optimized for STR genotyping. The curve for freebayes run without optimization is not shown due to poor performance.

We benchmarked HipSTR’s accuracy by comparing its STR calls from whole genome sequencing (WGS) data to capillary electrophoresis data, the current gold standard for STR genotyping. To this end, we obtained 118 WGS datasets from the Simons Genome Diversity Project (SGDP)^13^ that were sequenced with an Illumina 100bp pair-end PCR-free protocol to at least 30x coverage. These samples also have capillary electrophoresis calls for 600 highly polymorphic STRs from the Marshfield panel^14^, providing a challenging test case for STR callers. The capillary calls showed an internal consistency of 98.8% when comparing STR genotypes of a few duplicated samples, setting an upper bound on the accuracy that could be achieved in our tests. In addition to using HipSTR, we also genotyped the same exact STRs with three general-purpose callers and two STR-specific tools: GATK HaplotypeCaller (GATK-HC)^15^, Platypus^16^, freebayes^17^, lobSTR^5^ and RepeatSeq^6^. We optimized the set of command line options to boost the accuracy of each tool (Supplementary Table 1) and developed a machine learning approach to find a combination of quality indicators for each tool to optimally filter STR calls (Supplementary Table 2).

Our results show that HipSTR outperformed all other tools (Figure 1; Supplementary Table 3). Without any quality-based filtering of the Marshfield panel calls, HipSTR achieved an accuracy of 95.2%, better than any other tested method. In contrast, the second best tool (GATK-HC) achieved an accuracy of 92.0% with default settings and an accuracy of 94.3% after considerable optimization, reporting 20% more erroneous STR genotypes than HipSTR. When filtering the 10% least confident genotypes, HipSTR’s accuracy jumped to 98.8% and again substantially outperformed the next best tool.

We also evaluated the ability of HipSTR to go beyond reporting length polymorphisms to full STR haplotypes. About half of the STRs in the genome display a repeat structure that includes short interruptions to the recurrent motif^11^. Thus, two STR alleles that are identical by length can be discordant in sequence due to distinct evolutionary paths^18^ (Supplementary Fig. 5). Current STR callers and capillary electrophoresis methods only report an STR’s length and cannot differentiate between these homoplastic alleles. This complicates population genetics analyses, reduces the ability to understand the effect of STR variations on regulatory or coding changes, and limits the information available for forensic fingerprinting. As HipSTR automatically reports the full sequence of STR variants, not just their length, we sought to test the accuracy of our strategy by running it on the well-characterized CEPH trio in the Illumina Platinum genomes dataset. We observed ~70,700 STR loci that passed our filters in which at least two alleles had the same length but different sequence. Only 304 loci (0.4%) did not follow Mendelian consistency, showing the robustness of our method to report STR sequence variations. Next, for the same trio, we measured the ability of HipSTR to physically phase the Marshfield STR genotypes onto SNP haplotypes. For 178 loci where the algorithm confidently phased the child, we were also able to determine the transmitted paternal and maternal STR alleles. In all 178 instances, HipSTR correctly phased the paternal STR allele onto the paternal SNP scaffold, demonstrating the accuracy of its physical phasing approach. Taken together, our results highlight that HipSTR not only accurately reports STR length polymorphisms but also adds valuable information about the repeat structure and the haplotype context of the variants.

We found that HipSTR is scalable and apt to the analysis of large-scale sequencing data. We ran HipSTR on 2,000 Illumina whole genome sequencing datasets with at least 30x coverage from a New York Genome Center internal collection of genomes. We instructed HipSTR to genotype each of the 1.6 million STRs in the human reference genome. Analyzing this entire dataset in batches of 200 samples only required an average of 10 CPU hours per sample and at most 5GB of memory per batch. We were able to call STRs in this collection after a total of five days on the NYGC computing cluster. For each genome, HipSTR reported an average of ~360,000 STR loci that differed from the human reference.

Encouraged by the accuracy and scalability of HipSTR, we wondered about its ability to identity *de novo* STR mutations (Figure 2A). These mutations were not included in previous scans for *de novo* variants. After analyzing ~1.6 million STR loci in the CEPH trio, we detected ~745,000 loci with at least one length variation. To enhance the specificity of our analysis, we applied stringent quality filters and restricted our search for *de novo* mutations to ~265,000 STRs. Overall, HipSTR identified 423 *de novo* STR variants in which the child possessed an STR allele whose length was not in the parental genotypes.

**Figure 2:**
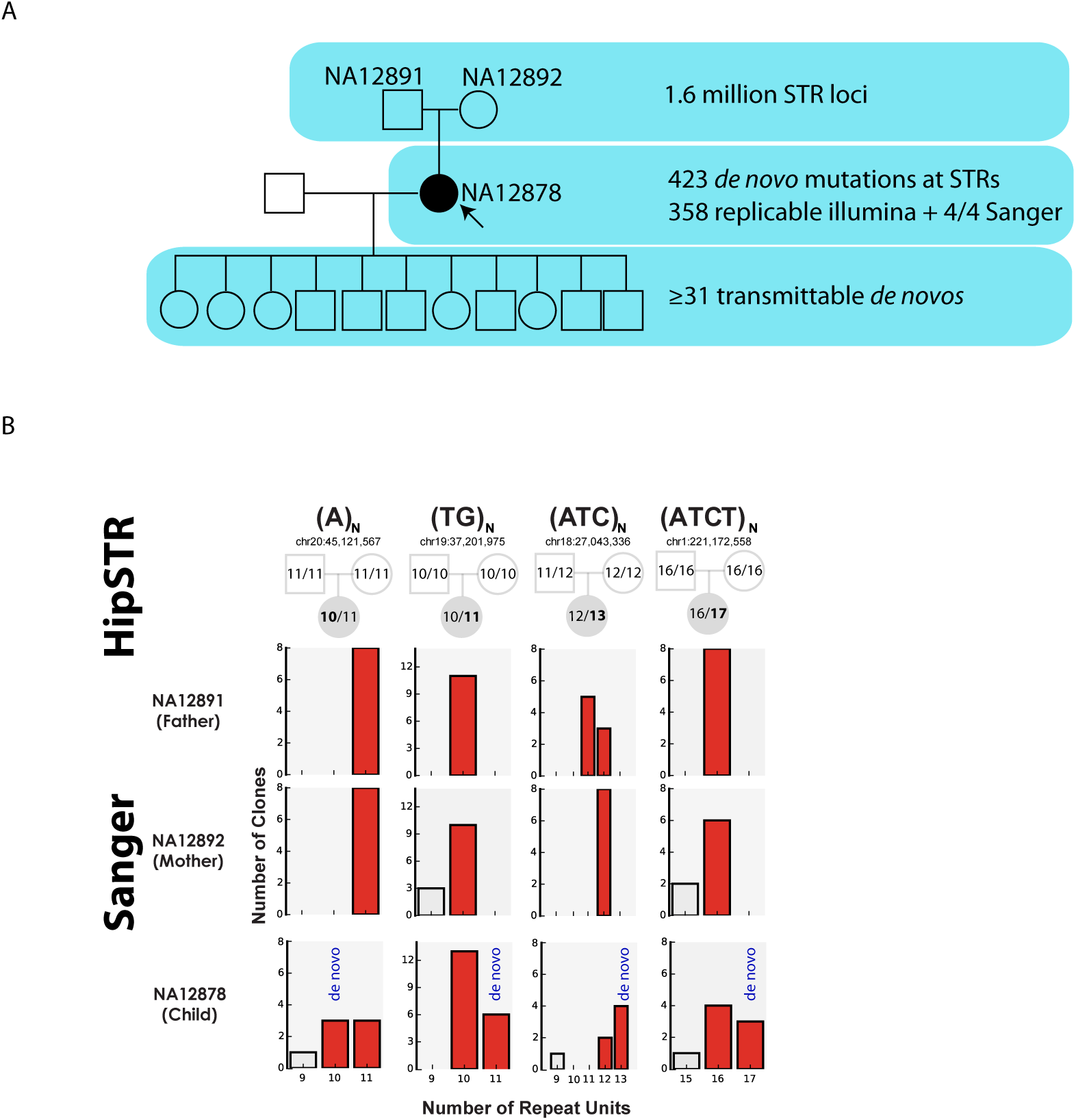
Genome-wide analysis of *de novo* STR mutations. **(A)** The experimental stages of identifying *de novo* mutations in NA12878 **(B)** Validating *de novo* STR mutations. Top: the number of repeats observed by HipSTR in the trio (bold: *de novo* allele). Bottom: the Sanger results for each individual. In all instances, we observed multiple clones supporting the *de novo* allele in NA12878 but not in the parents. Multiple clones also supported each of the alleles predicted by HipSTR (red bars), whereas clones containing unpredicted alleles (gray bars) likely due to stutter were observed with much lower frequency.

To validate these mutations, we re-ran HipSTR with distinct Illumina datasets generated for these samples and compared the variants between the runs. Notably, we were able replicate 358 (85%) of the *de novo* mutations as all samples’ allele lengths matched perfectly between the two call sets. As expected, most *de novo* mutations predominantly occur at homopolymer repeats (293/358=81%) and to a much lesser extent at repeats with 2-6 base pair repeat units. To further validate our results, we genotyped high confidence *de novo* STR loci with capillary electrophoresis. As these STRs are highly mutable and prone to stutter noise, even Sanger sequencing does not deliver unequivocal genotypes for these loci. To gain confidence, we TOPO cloned the STR alleles from each member of the trio and Sanger sequenced at least eight independent clones per individual. Using this technique, we obtained high confidence Sanger calls for four STRs where HipSTR identified a *de novo* mutation (Figure 2B, Supplementary Figs. 6-9). In all cases, the Sanger calls for the parents confirmed the parental genotypes reported by HipSTR and the Sanger calls for the child confirmed the *de novo* allele reported by HipSTR, validating our method.

Next, we sought to distinguish *de novo* STR variants that arose in the germline of the trio’s child (NA12878) from mutations that arose during the many cell-line passages of this sample. We used HipSTR to analyze the WGS data of her 11 offspring and inspected the genotypes for the 358 STRs where NA12878 had a replicable *de novo* mutation. For 31 loci, we found evidence that the *de novo* STR allele was transmitted to at least three offspring and was absent from the paternal genotype. For an additional 32 loci, the *de novo* allele of the mother was observed in at least three of her offspring, but the same (identical-by-state) allele also resided in her husband (NA12877) and the transmission could not be fully resolved. As our genome-wide scan identified at least 31 *de novo* STR mutations in the germline of NA12878 after examining ~35% of STRs in her genome, we predict a lower bound of over 100 *de novo* STR mutations per generation. Previous studies of *de novo* SNP rates quantified approximately 70 mutations per generation^8, 19^. Our analysis strongly suggests that the load of STR *de novo* variations is at least of the same order of magnitude, demonstrating their substantial source of genetic variability.

To summarize, our results show that HipSTR offers several advantages for STR calling. First, the technique is considerably more accurate than other general purpose and STR-specific variant callers and does so with exceptional computational tractability. Second, the tool has new capabilities such as phasing, haplotyping, and resolving STR homoplasy that are absent from other tools and can be important for population genetic analyses, forensic work, and STR association studies. Finally, our method enables highly specific detection of *de novo* STR variants. This new capability could open exciting avenues for understanding human genetic variation and the role of STRs in various diseases and complex traits.

## Methods

### The HipSTR algorithm

#### Modeling PCR stutter

PCR stutter artifacts add or remove copies of an STR’s motif to sequencing reads, resulting in observed STRs that differ in size from the true underlying genotype. To mitigate these effects, we used a model that we developed and extensively validated in our previous work to discern between stutter noise and STR mutations on the Y chromosome^12^. HipSTR constructs a stutter model *θ*_*x*_ for each STR locus *x*, which contains the probability that stutter adds (*u*) or removes (*d*) repeats from the true allele in an observed read, and a geometric distribution with parameter *ρ*_*s*_ that controls the size of the stutter-induced changes. In our framework, the probability of observing a stutter artifact δ repeat units in size is:

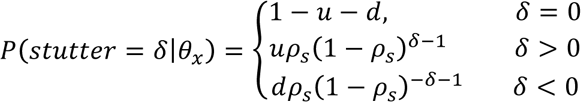

To estimate each locus’ stutter model parameters, we extract the size of the STR observed in each read for all individuals in the population. We then use an Expectation-Maximization approach^20^ to learn the parameters. The E-step computes each sample’s genotype posteriors:

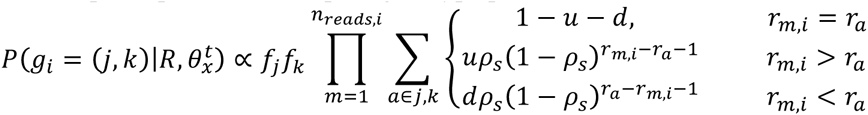

Here, *R* denotes the set of all reads, *g*_*i*_ denotes the phased genotype for the *i*^*th*^ individual, *n*_*reads*,*i*_ denotes the number of reads for the *i*^*th*^ individual, r_m,*i*_ denotes the number of repeats in the *m*^*th*^ read for the *i*^*th*^ individual, r_*a*_ denotes the number of repeats in the *a*^*th*^ allele and f_*j*_ denotes the frequency of the *j*^*th*^ allele. For each possible phased genotype, the E-step also computes the conditional probability that a read originated from either allele:

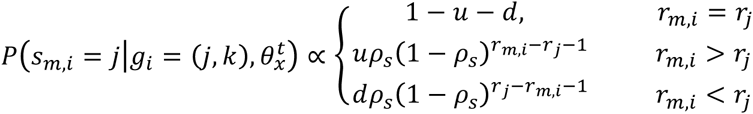

Given *N* samples, *A* alleles and *Q* reads, the M-step then updates the stutter model parameters and allele frequencies using these probabilities:

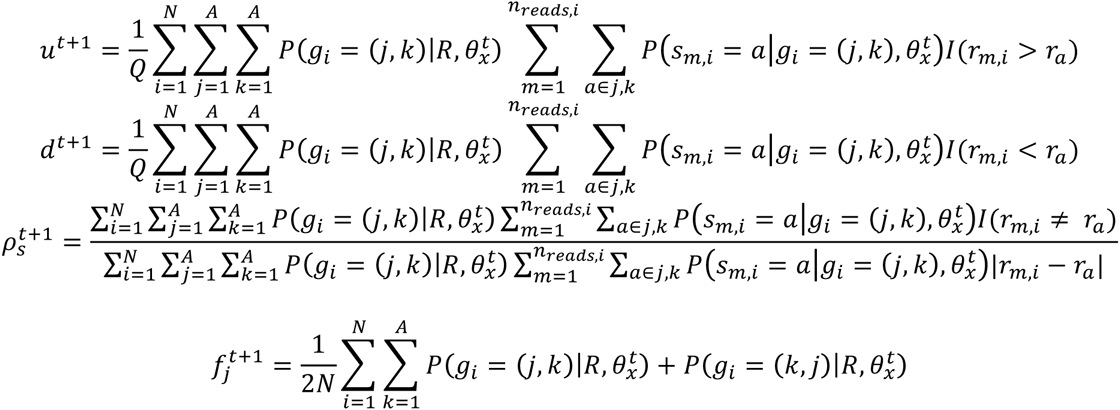

Intuitively, the update rules for the stutter probabilities *u* and *d* compute the fraction of times a read’s STR allele is either larger or smaller than its underlying allele. The update rule for the step size parameter *ρ*_*s*_ is more involved, but it first restricts the computation to reads with non-zero stutter. It then computes the inverse of the mean weighted step size, consistent with a maximum likelihood estimator for a geometric distribution.

#### Generating candidate alleles

To identify an initial set of STR alleles, HipSTR selects reads that fully span the STR. It requires that both ends of a read match the reference genome for at least 10bp and that neither end of the read has a longer exact match with the reference genome 15bp upstream or downstream from its alignment. Based on this subset of reads, HipSTR includes a sequence as a candidate allele if is present in two or more and at least 20% of a sample’s reads.

HipSTR also uses an iterative approach to identify new candidate alleles. At the end of every round of genotyping, it computes the maximum-likelihood genotype for each sample and realigns every read relative to the most probable allele in its sample’s genotype. Each of these alignments generates a sequence in the STR region. If the same sequence is observed in a sample in two or more alignments with stutter artifacts, HipSTR selects the sequence as a new candidate allele.

#### Computing genotype likelihoods

The genotype likelihood model integrates information about every read’s phasing likelihood and alignment likelihood. For the *m*^*th*^ read for individual *i*, *P*(*p*_*m*,*i*_|h = 1) and *P*(*p*_*m*,*i*_|*h* = 2) denote the phasing likelihoods of the read originating from the first and second SNP haplotypes, while *P*(*s*_*m*,*i*_ | *a* = *j*) denotes the alignment likelihood of the read to the *j*^*th*^ allele. We use a uniform prior for each unphased genotype, such that heterozygous phased genotypes have half the prior probability of their homozygous counterparts. The likelihoods for the phased genotypes for the *i*^*th*^ sample are:

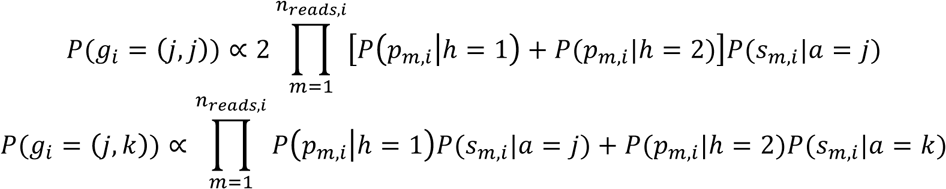

#### Read phasing likelihoods

To compute the phasing likelihoods for each read, HipSTR examines bases in the read or its mate pair that are aligned to heterozygous SNPs. If the read originated from a haplotype, the likelihood of the base *b*_*i*_ matching the SNP base h_*j*_ is given by the base quality *q*_*bi*_ while the likelihood of it not matching is one third of the residual probability. We express this as:

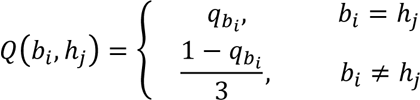

We compute each read’s total phasing likelihood by multiplying *Q*(*b*_*i*_, *h*_*j*_) for every base *b*_*i*_ aligned to a heterozygous haplotype SNP *h*_*j*_ in the read or its mate pair. In practice, SNP calls in and around STR regions are likely to be error-prone. We therefore exclude SNPs that are within 15 base pairs of the STR region when computing the phasing likelihoods.

#### Read alignment likelihoods

HipSTR assumes that each haplotype is composed of two distinct types of regions: flanking sequences and STR sequences. As the sources of error prevalent in these two types of regions differ dramatically, HipSTR uses a distinct model to align sequences to each type of region. It then combines the likelihoods from these two different models at the junctions of these regions. In particular, it requires that the read match the flanking sequence at the first base preceding the STR sequence and at the first base following an STR sequence.

#### Aligning reads to flanking sequences

Within flanking sequences, the alignment model accounts for Illumina sequencing errors using a previously described hidden Markov model^21^. To efficiently align reads in these regions, we use three matrices to recursively compute the maximum log-likelihood of aligning read bases *b*_l_… *b*_*i*_ with haplotype bases *h*_1_… *h*_*j*_. The matrices *Match* and *Ins* are used to track the likelihoods that read base *b*_*i*_ is aligned to haplotype base *h*_*j*_ or an insertion following base *h*_*j*_, respectively. Matrix *Del* tracks the maximum log-likelihood that base *b*_*i*_ is followed by one or more deletions. In conjunction with values for *t*_*X* ➔ *Y*_, the log-probability of transitioning from hidden state *X* to hidden state *Y*, we use the following recursions to fill in each matrix column-by-column:

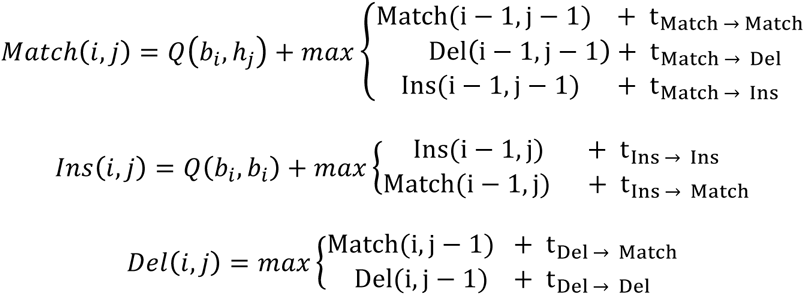

#### Aligning reads to STR sequences

In STR regions, HipSTR utilizes an entirely different alignment model to account for STR-specific errors. As PCR stutter artifacts are prevalent in this domain, it assumes that a read’s sequence differs from the underlying haplotype by at most one indel whose magnitude *D* is a multiple of the repeat unit length. If no stutter artifact has occurred, the likelihood of the observed sequence is governed by the agreement between each base in the read and its corresponding haplotype base. The probability of no stutter artifact and aligning base *b*_*i*_ and its preceding bases to an STR sequence *h*_1_… *h*_*L*_ of length *L* is:

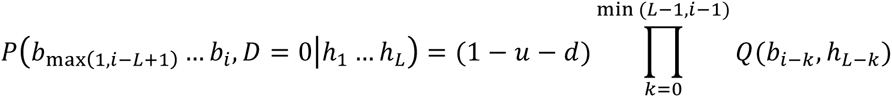

If a stutter deletion occurs, we assume that it can arise anywhere within the STR region. We iterate over these configurations, each of which has a likelihood given by the agreement between the sequenced bases and their corresponding haplotype bases:

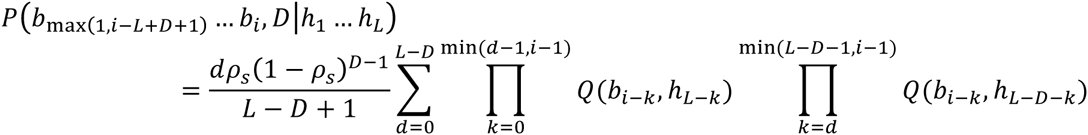

Finally, if a stutter insertion occurs, we assume that it can precede any base in the STR region. As PCR stutter insertions typically contain sequences that copy the local repeat structure, we assume that inserted sequences are periodic copies of the STR sequence directly preceding the insertion. We therefore measure the likelihood of inserted bases according to their agreement with this sequence. For an STR with a repeat motif of length *M*, iterating over each possible insertion position results in the likelihood:

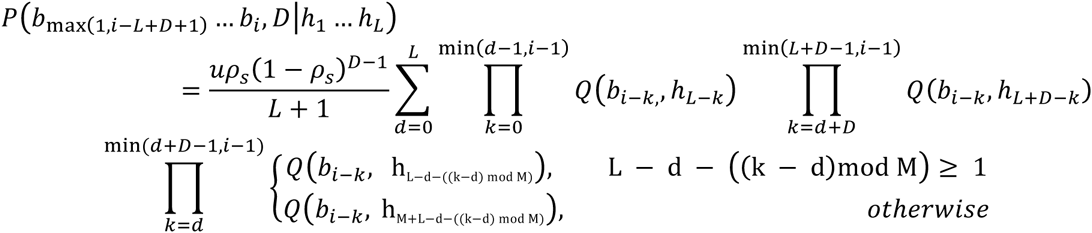

## Experiments

### Constructing a gold standard STR dataset

We downloaded capillary genotypes for the 628 STRs of the Marshfield panel from <u>https://web.stanford.edu/group/rosenberglab/data/rosenbergEtAl2005/combinedmicrosats-1048.stru</u> and other information from <u>https://web.stanford.edu/group/rosenberglab/data/pembertonEtAl2009/Pemberton_AdditionalFile1_11242009.txt</u>. These datasets were generated in a prior study^14^. Using the isPCR^22^ tool, we mapped each STR’s primers to the hg19 reference genome. Together with available repeat structure information, we used Tandem Repeats Finder^23^ to scan between the primer sites and pinpoint the exact genomic coordinates of each STR.

Capillary STR genotypes report the lengths of amplified DNA fragments containing the STR region. These lengths correctly capture STR variations but also capture indels outside of STRs if they fall within the amplified regions. To mitigate the effects of these indels, we downloaded assembly-based FermiKit^24^ calls for each of the 263 SGDP samples from a publicly available repository (https://github.com/lh3/sgdp-fermi). For each STR, we identified samples with indels located in between the two primer sites. We then masked the genotype for a sample if any of its indels occurred more than 15bp upstream or downstream from the STR region, as these are unlikely to originate from the STR itself.

#### Assessing variant caller performance

We downloaded BWA-MEM alignments^25^ available for 263 publicly available SGDP samples (accession ERP010710). Using a BED file containing each STR’s genomic coordinates, we ran every tool with default options. We then explored the effects of alternate command line options by rerunning each tool using every combination of the settings listed in Supplementary Table 1. The combination of settings that led to the highest level of agreement with the capillary genotypes was then selected as optimal. Despite multiple attempts, we were unable to run STR-FM^7^ outside of the Galaxy environment and therefore could not include this tool in our benchmarking experiments.

Most general-purpose variant callers produce multiple unphased variants per STR region. Without phase information, it is frequently not possible to determine the lengths of an individual’s two STR alleles present at a given locus. To overcome this issue, we summed the sizes of all indels each caller produced within an STR region for a given sample. The frequency with which these sums exactly matched those predicted by the capillary electrophoresis data was then used as the variant caller’s accuracy.

#### Phased trio SNP scaffolds

We used the HaplotypeCaller module in GATK v3.5-0-g36282e4 to jointly genotype all members of the trio using the aligned and sorted BAMs for runs ERR194147, ERR194160 and ERR194161 generated as described above. In accordance with guidelines for hard filtering SNP calls, we used GATK’s SelectVariants and VariantFiltration modules to select only those SNPs with a passing FILTER, *QD* > 2, *FS* < 60, *MQ* > 40, *MQRankSum* > -12.5 and *ReadPosRankSum* > -8. Next, we downloaded v5a of Beagle’s^26^ reference panels for Phase 3 of the 1000 Genomes Project from the tool’s website and removed three samples that are part of the CEPH pedigree (NA12878, NA12889 and NA12890). Using v4.0-r1399 of Beagle and these filtered reference panels, we phased the trio’s SNP calls using the *ped* option and *phase-its*=10, *burnin-its*=10, *impute-its*=0 and *impute*=false.

#### Generating CEPH trio STR genotypes for the Marshfield markers

We downloaded FASTQ files from the Illumina Platinum Genomes Project and an additional study of the effects of PCR amplification on sequencing errors^27^, resulting in data from twelve different sequencing runs (NA12878: ERR194147, SRR826463, SRR826467, SRR826469; NA12891: ERR194160, SRR826427, SRR826448, SRR826465; NA12892: ERR194161, SRR826428, SRR826473, SRR826471). We individually aligned each run to the hg19 reference genome using the BWA-MEM algorithm from bwa v0.7.12-r1039. Using the resulting BAMs for all samples concurrently, we ran HipSTRv0.2 with the options *--def-stutter-model --use-all-reads --min-reads* 25 and *--read-qual-trim #*. To generate STR genotypes that are phased onto SNP scaffolds, we reran HipSTR using the same arguments but also specified the *--snp-vcf* option and used the VCF containing phased SNP scaffolds as input.

#### Evaluating HipSTR’s physical phasing accuracy

We began by restricting our analysis to Marshfield markers in which the STR alleles transmitted from each parent to the child could be determined from the unphased genotypes alone. We then required that the child have a confidently phased STR genotype as indicated by a FORMAT field with PQ > 0.9 (minimum phased genotype posterior of 90%). Using the SNPs 50 kilobases upstream and 50 kilobases downstream of the STR region, we determined the SNP haplotype each parent transmitted to the child by requiring that each of the child’s SNP haplotypes exactly match one parental haplotype and that these matches involve both the maternal and paternal haplotypes. After enforcing each of these requirements, 178 markers were available for downstream analyses. For each of these markers, HipSTR’s phased genotypes correctly placed the paternally transmitted STR allele onto the paternally transmitted SNP haplotype, resulting in perfect phasing accuracy.

#### Running HipSTR on a population-scale dataset

WGS data for 2000 individuals sequenced using 150bp paired-end Illumina reads and more than 30x coverage was internally available at the New York Genome Center. We applied HipSTR to genotype STRs in this dataset. Using the BWA-MEM aligned BAMs for each sample as input, we jointly genotyped 200 samples at a time with HipSTR and all default options. We aggregated the timing statistics across each of the individual runs to determine the total run time of ~20,000 CPU hours. NYGC’s IRB committee approved all human subject experiments prior to this analysis.

#### Using HipSTR to identify *de novo* mutations

We downloaded FASTQs from the Illumina Platinum Genomes project containing 200x sequencing data for NA12877 and NA12878 (runs ERR174310-ERR174341). As before, we used BWA-MEM to align the reads in each of these runs individually. Using all of these BAMs and two previously generated BAMs for NA12891 and NA12892 (ERR194160 and ERR194161), we ran HipSTR with the options *-- def-stutter-model –require-pairs, --min-reads* 25 and a BED file containing 1.6 million STR regions.

We applied a series of stringent filters to the call set to reduce the likelihood that genotyping errors introduce false positive *de novo* calls. Using the FORMAT fields available in the HipSTR VCF, we required that all three individuals in the trio have a minimum genotype posterior (Q) of 0.9, no more than 10% of reads containing either a stutter artifact or a flanking sequence indel (DSTUTTER/DP and DFLANKINDEL/DP) and at least 10 reads spanning the STR region (MALLREADS). Lastly, we required that the ratio of spanning reads supporting the alleles for each individual be at least 20% (MALLREADS).

To identify an initial set of candidate mutations, we examined STRs that satisfied all of these requirements. For 423 loci where the child (NA12878) carried an allele length not observed in either of the parents (N12891 and NA12892), we identified a potential *de novo* mutation.

#### Validating *de novo* mutations using orthogonal datasets

We downloaded FASTQs containing 300x Illumina sequencing data for NA12878 from the Genome in a Bottle consortium and FASTQs for NA12878, NA12891 and NA12891 from the 1000 Genomes Project (SRR622457, SRR622458 and SRR622459, respectively). After generating BAM files using BWA-MEM, we collated these alignments with others generated in previous analyses (NA12878: SRR826463, SRR826467, SRR826469; NA12891: SRR826427, SRR826448, SRR826465; NA12892: SRR826428, SRR826471, SRR826473). We then reran HipSTR using these BAMs and the options *--def-stutter-model--require-pairs --min-reads* 25 and a BED file containing the 423 STR regions with previously detected *de novo* mutations.

Without performing any filtering, we compared the HipSTR calls from these datasets to the calls generated during the discovery phase. For 358 of the markers, each member of the trio had allele lengths that matched perfectly between the two call sets, resulting in a set of sites with high confidence *de novo* mutations. The genotypes for these STRs with high confidence *de novo* mutation are available as a VCF at https://github.com/HipSTR-Tool/HipSTR-paper.

#### Sanger sequencing validation

Primers were designed around the STR coordinates to generate 300-600 bp PCR products (Supplementary Table 4). All coordinates are according to GRCh37/hg19. Primers were tested using *in silico PCR* for unique products. Genomic DNA for NA12878, NA12891, and NA12892 was obtained from the Coriell Institute (Camden, NJ, USA). DNA was amplified for 30 cycles in 25 ul reactions according to the manufacturer’s recommended cycling conditions using Q5 High Fidelity Polymerase (NEB catalog #M0494) to reduce stutter, generating blunt end products. Amplicons were purified on magnetic beads (Thermo Fisher Scientific ChargeSwitch PCR Clean-Up Kit, catalog #CS12000) and cloned into linearized pMiniT (NEB catalog #E1202). Plasmids were transformed into 50ul of chemically competent E. coli (Lucigen E. cloni Chemically Competent Cells catalog #60108). Outgrowth cultures (50ul) were incubated overnight on ampicillin plates. Individual colonies were selected and cultured overnight in 2mL LB + ampicillin (100ug/mL). DNA was extracted and column purified (Thermo Fisher Scientific PureLink Quick Plasmid Miniprep Kit, catalog #K210010). Sanger sequencing of at least 8 clones per individual per locus was performed by Eton Bioscience (Newark, NJ, USA) using the supplied primers for the pMiniT plasmid. Only results with flanking sequences upstream and downstream of the STR of sufficient quality were included in the final counts.

#### Assessing *de novo* transmission to children

We downloaded FASTQs containing 50x Illumina sequencing data for each of the 11 children of NA12878 (ERR194148, ERR218433, ERR324432, ERR324433, ERR194152, ERR324434, ERR194154, ERR194155, ERR324435, ERR194157 and ERR194162). Using these BAMs as input, we ran HipSTR with the options *--def-stutter-model --require-pairs --min-reads* 25.

#### Coordinates

All reported coordinates are based on the hg19 genome build.

## Acknowledgments

Y.E. holds a Career Award at the Scientific Interface from the Burroughs Wellcome Fund. This study was supported by NIJ grant 2014-DN-BX-K089 (T.W., D.Z, A.G., M.G., Y.E.) and a generous gift by Andria and Paul Heafy.

---

**Supplementary Figure 1:**
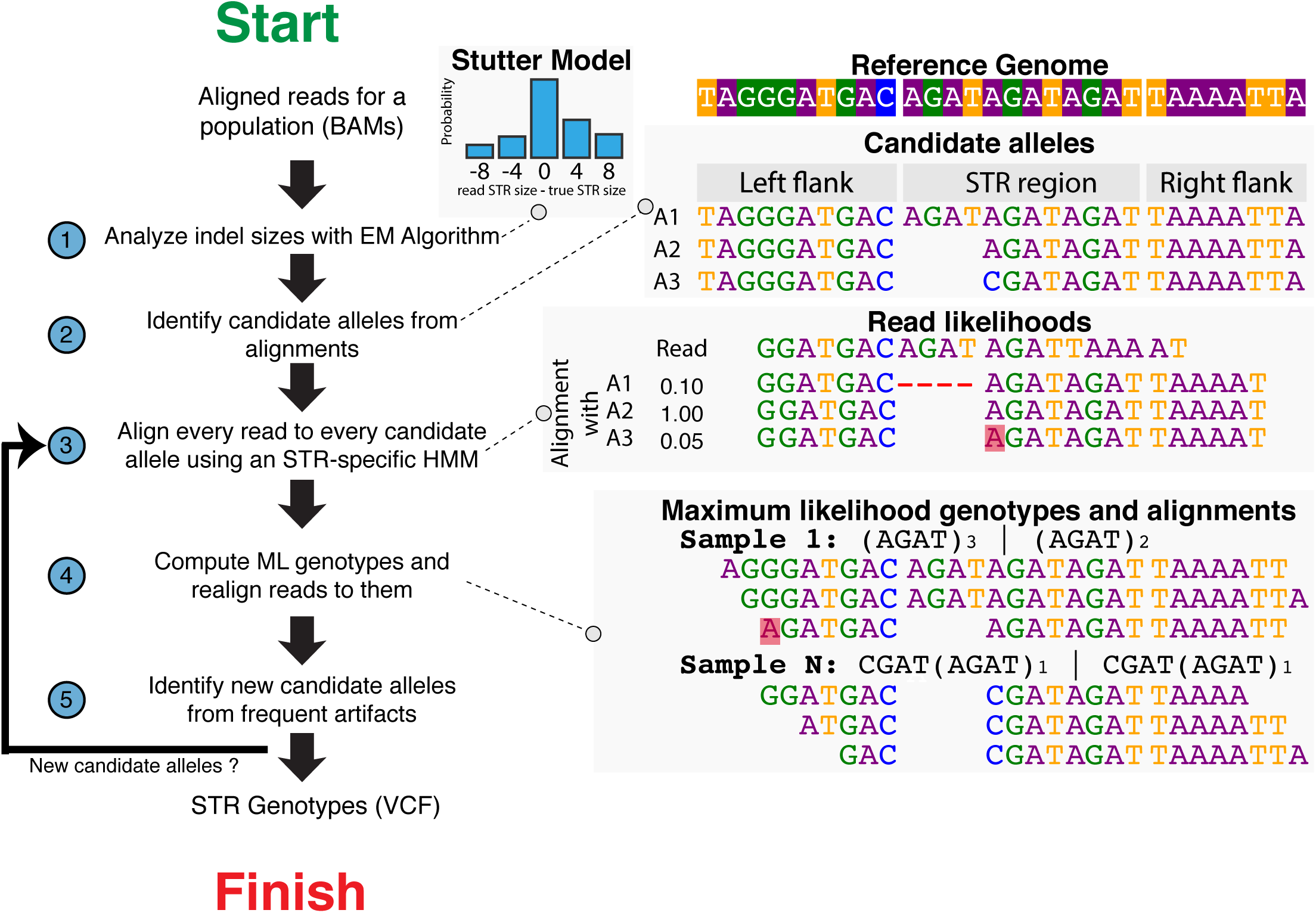
Overview of the HipSTR algorithm. In step 1, an expectation-maximization algorithm learns the PCR stutter model for the locus of interest in order to account for the frequency of these errors during genotyping. Step 2 utilizes well-anchored alignments that span the STR to identify candidate alleles in the STR region and builds haplotypes using these alleles and the sequence upstream and downstream of the STR. In step 3, the PCR stutter model and an HMM are used to align every read to every candidate haplotype. Step 4 analyzes reads that overlap heterozygous SNPs to determine the likelihood that each read came from either strand of the phased SNP haplotypes. Finally, step 5 combines these two sets of likelihoods to determine each sample’s maximum likelihood genotype. Every read is realigned relative to its sample’s optimal genotype and if new reads span the STR, HipSTR returns to step 2 to identify new candidate alleles and repeat the process. This iterative procedure continues until no new candidate alleles are identified, at which point the maximum likelihood genotypes are output to a VCF file.

**Supplementary Figure 2:**
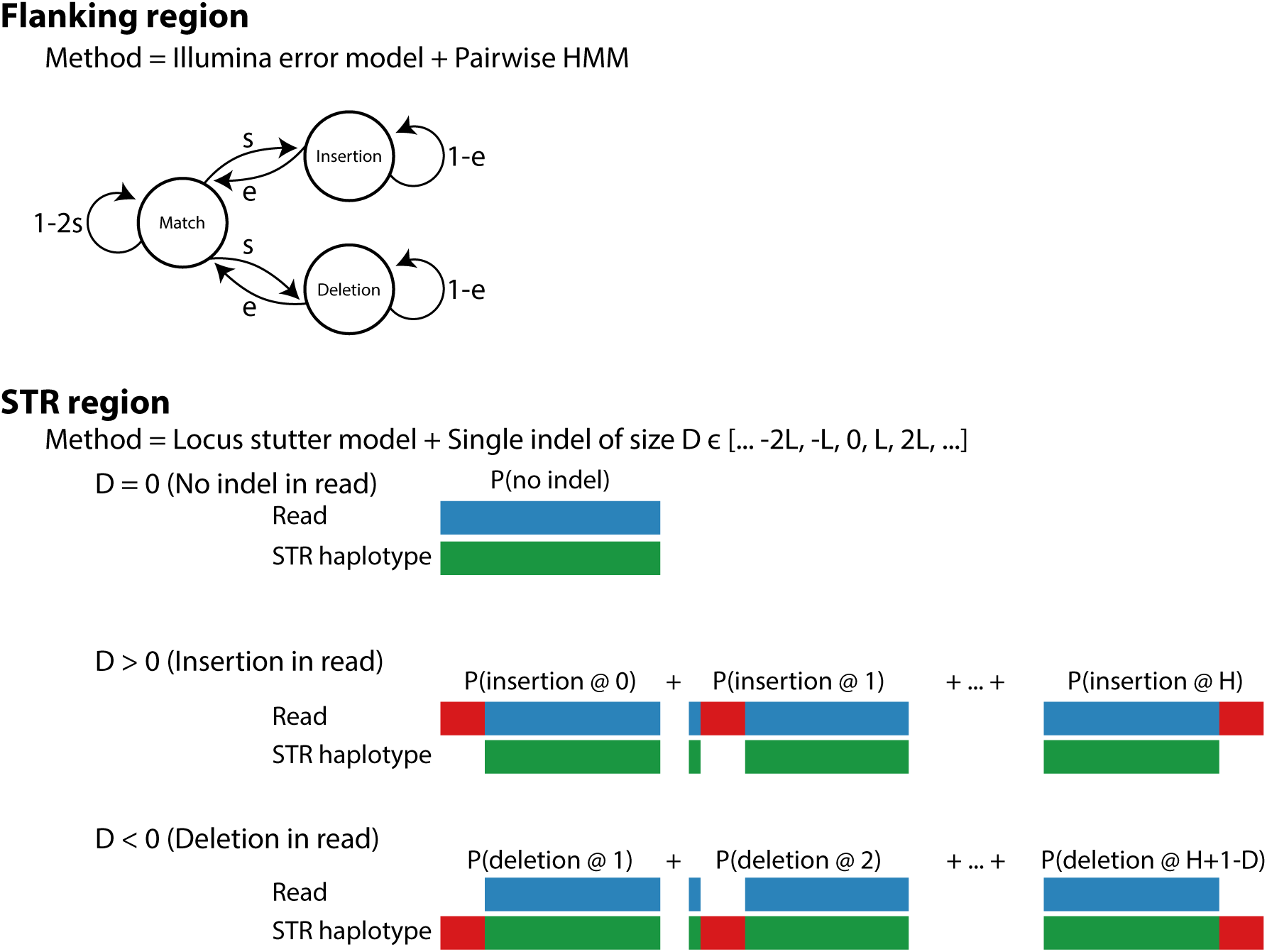
HipSTR alignment models. HipSTR uses two distinct types of alignment models to obtain robust alignments likelihoods. In regions flanking the STR (top), a pairwise hidden Markov model is used to accounts for Illumina sequencing errors. In contrast, within STR regions (bottom), HipSTR assumes that the main source of artifacts are stutter errors. If no stutter error occurs, the likelihood of observing a sequence of characters is given by the agreement between the bases in the read (blue) and the corresponding bases on the haplotype (green). Otherwise, HipSTR assumes that a single stutter indel occurs (red) and that it arose at each position with equal probability. The alignment likelihood for these scenarios are then obtained by marginalizing over all configurations.

**Supplementary Figure 3:**
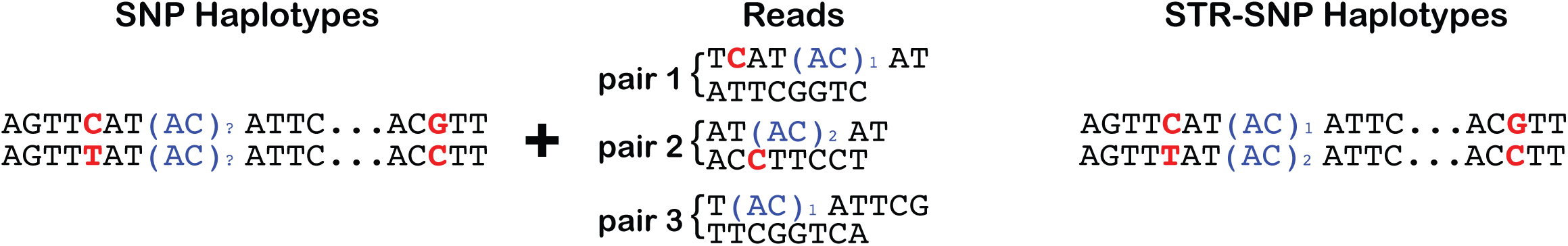
Physically phasing STRs onto SNP scaffolds. When provided with phased SNP haplotypes, HipSTR uses reads that overlap heterozygous SNPs to phase the STR genotypes (blue) onto the SNP haplotypes (red). The schematic provides a conceptual oultine of how this works for an AC repeat flanked by two heterozygous SNPs with known phase.

**Supplementary Figure 4:**
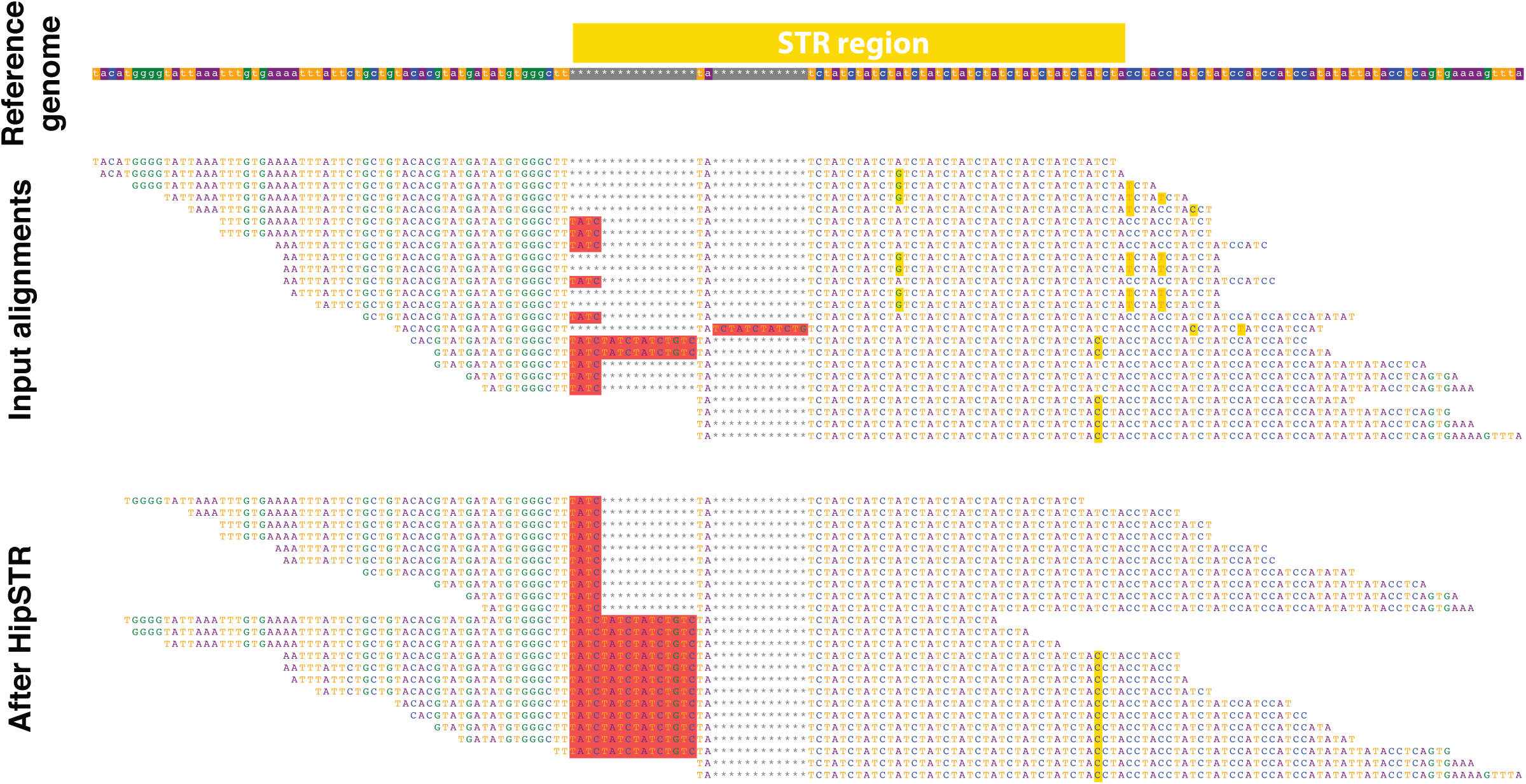
HipSTR in action. Example alignments for NA12878 to a particular Marshfield STR before (top) and after (bottom) processing by HipSTR. While the input alignments are riddled with local alignment errors, HipSTR’s algorithm resolves the alignments into two parsimonious STR insertions of 4 and 16 base pairs.

**Supplementary Figure 5:**
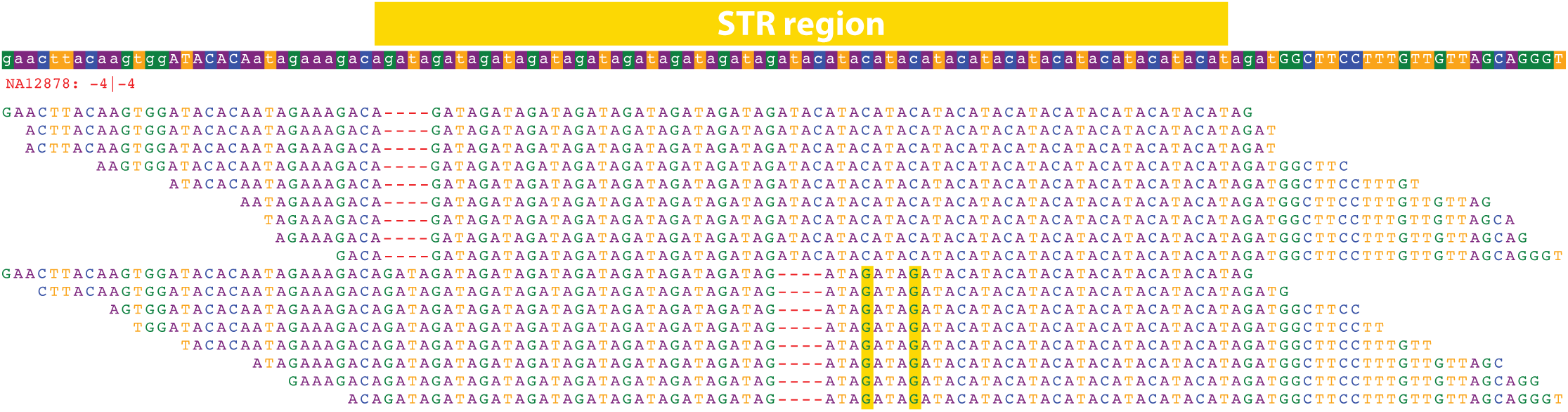
An example of STR homoplasy. The figure depicts an STR located at chr6:16429779 in the hg19 reference genome with an AGAT repeat followed by an ACAT repeat (top box). Each subsequent row depicts the maximum likelihood alignment for reads from sample NA12878 after processing with HipSTR. While all of the reads support a 4bp deletion, they support two different STR sequences. In the top 9 reads, one AGAT unit is deleted while the ACAT perfectly matches the reference. In the bottom 8 reads, two copies of AGAT are inserted followed by a deletion of three ACAT copies. HipSTR correctly disentanges these two haplotypes and reports genotypes of (AGAT)_8_ (ACAT)_9_ and (AGAT)_10_ (ACAT)_6_ for NA12878 at this locus.

**Supplementary Figure 6:**
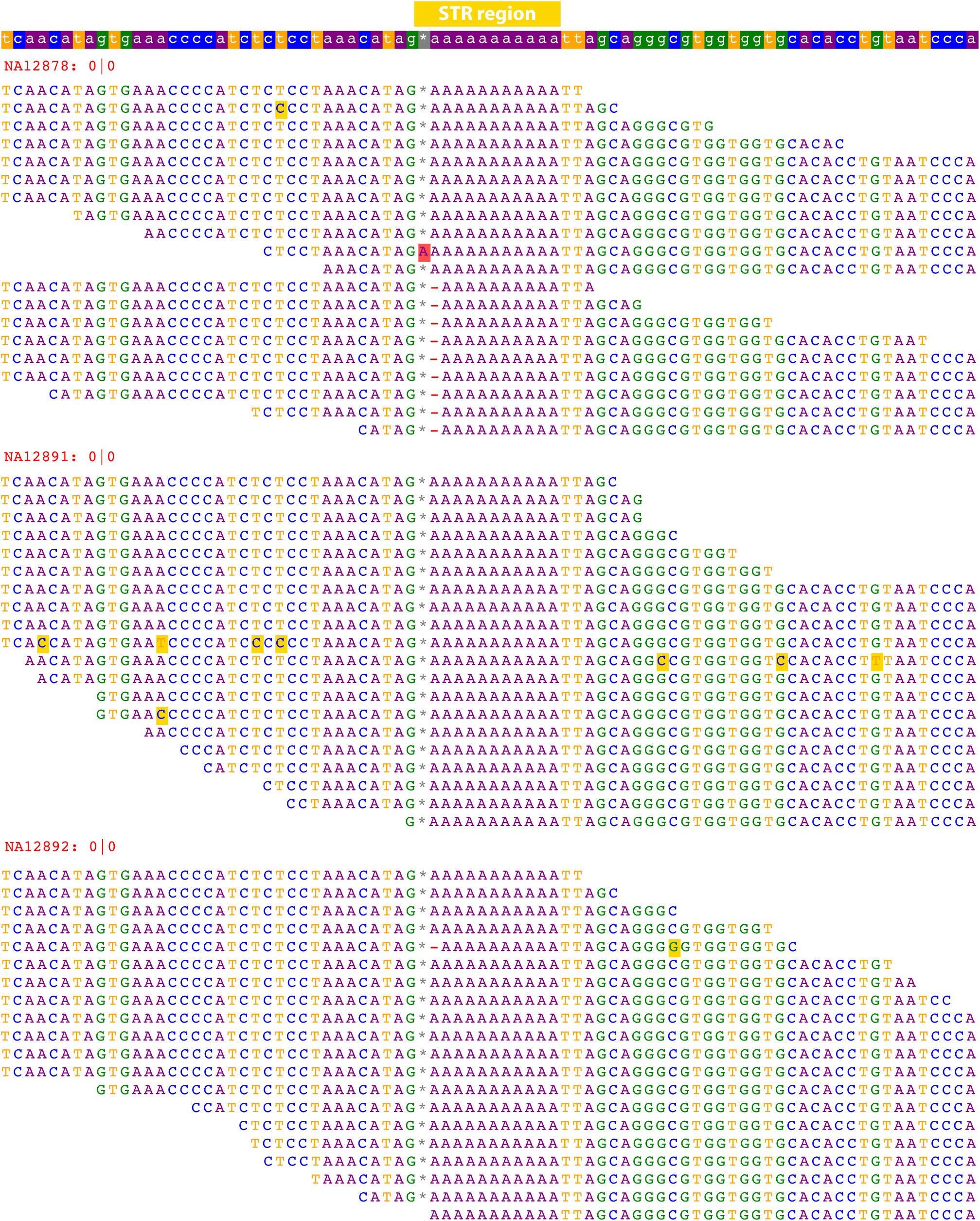
High-confidence de novo mutation detected by HipSTR. The figure depicts an STR located at chr20:45121567 in the hg19 reference genome with an A repeat (top row). Each subsequent row depicts a read’s maximum likelihood alignment after processing with HipSTR for the child (NA12878), father (NA12891) and mother (NA12892) in the trio. While the alignments strongly support that both the father and mother match the reference allele, the alignments for NA12878 support the reference allele and a 1bp deletion. Only a subset of the alignments for NA12878 (12.5%), NA12891 (50%) and NA12892 (50%) are displayed to facilitate visualization.

**Supplementary Figure 7:**
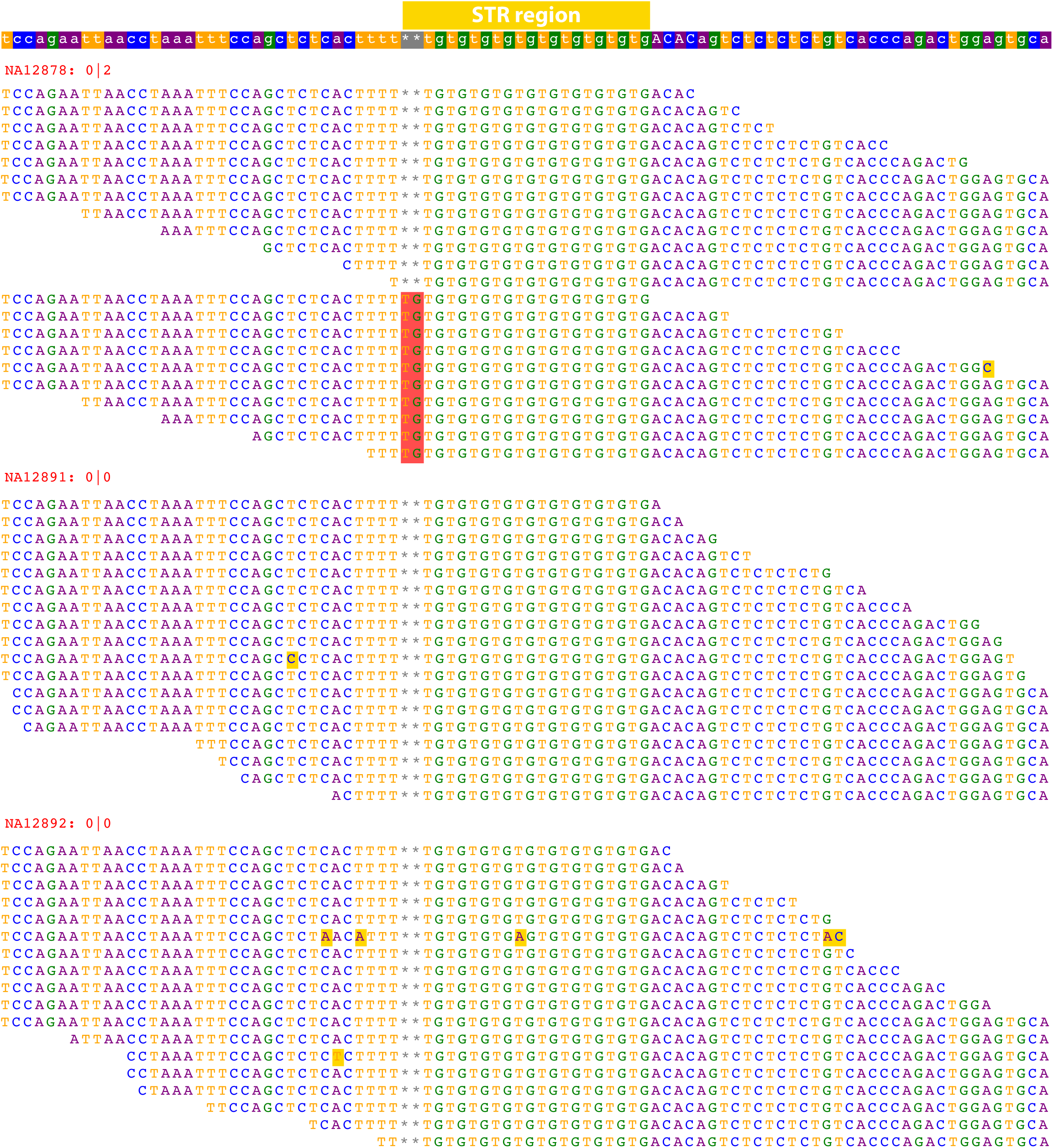
High-confidence de novo mutation detected by HipSTR. The figure depicts an STR located at chr19:37201975 in the hg19 reference genome with a TG repeat (top row). Each subsequent row depicts a read’s maximum likelihood alignment after processing with HipSTR for the child (NA12878), father (NA12891) and mother (NA12892) in the trio. While the alignments strongly support that both the father and mother match the reference allele, the alignments for NA12878 support both the reference allele and a 2bp insertion. Only a subset of the alignments for NA12878 (12.5%) and NA12892 (50%) are displayed to facilitate visualization.

**Supplementary Figure 8:**
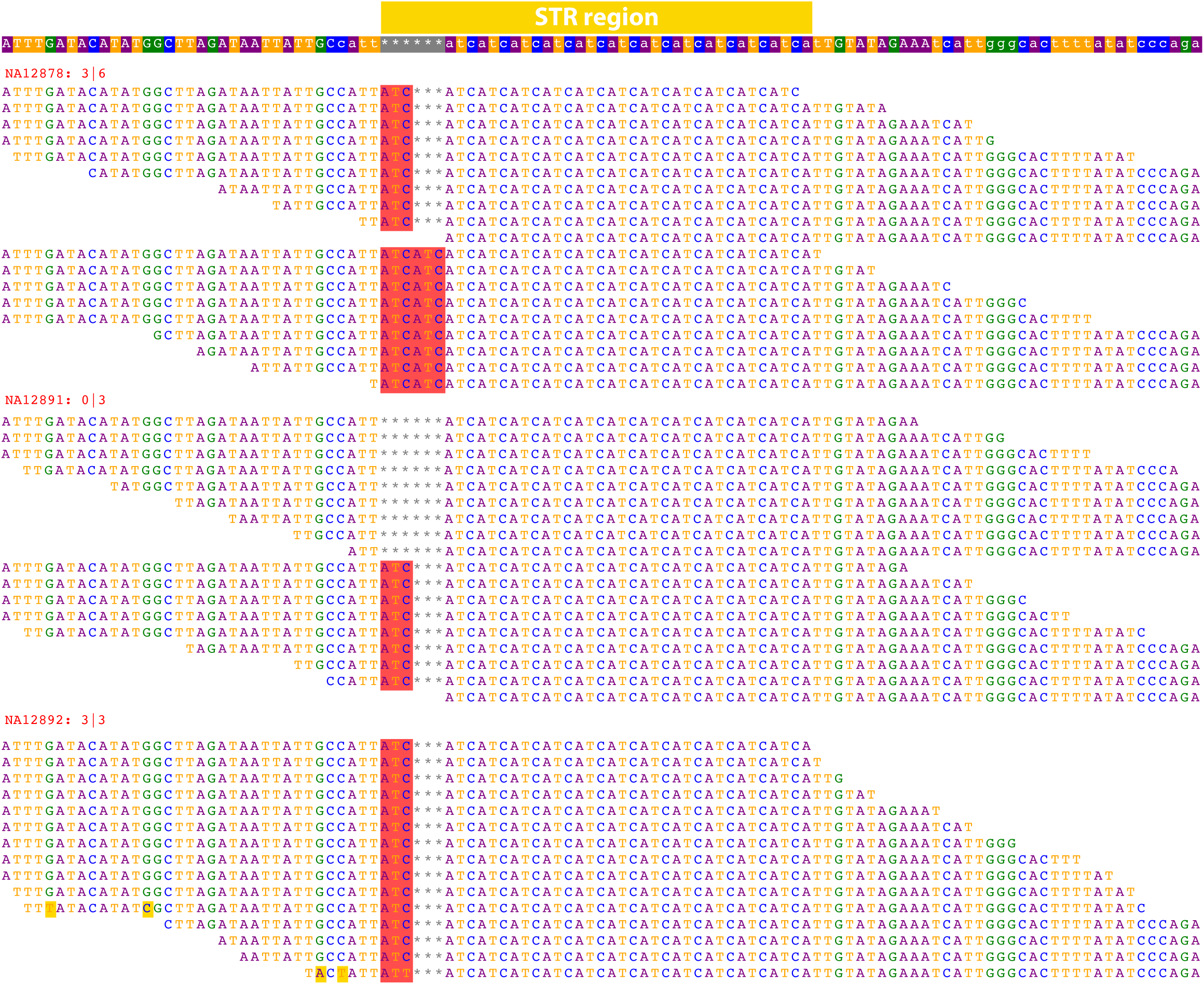
High-confidence de novo mutation detected by HipSTR. The figure depicts an STR located at chr18:27043336 in the hg19 reference genome with an ATC repeat (top row). Each subsequent row depicts a read’s maximum likelihood alignment after processing with HipSTR for the child (NA12878), father (NA12891) and mother (NA12892) in the trio. While the alignments strongly support a 0 bp/3bp heterozygous insertion for the father and a 3bp homozygous insertion for the mother, the alignments for NA12878 support a heterozygous 3bp/6bp insertion. Only a subset of the alignments for NA12878 (12.5%), NA12891 (50%) and NA12892 (50%) are displayed to facilitate

**Supplementary Figure 9:**
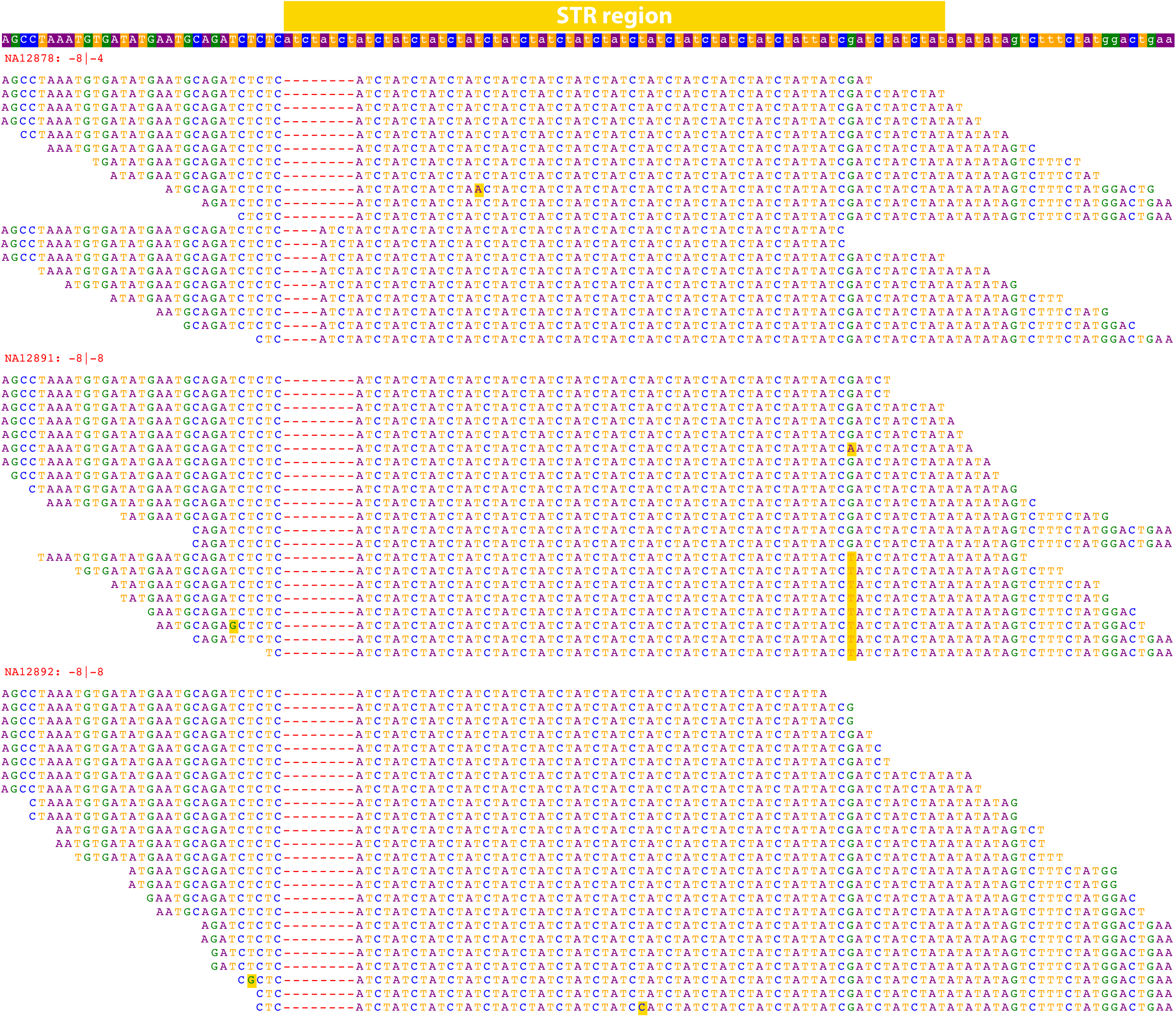
High-confidence de novo mutation detected by HipSTR. The figure depicts an STR located at chr1:221172558 in the hg19 reference genome with an ATCT repeat (top row). Each subsequent row depicts a read’s maximum likelihood alignment after processing with HipSTR for the child (NA12878), father (NA12891) and mother (NA12892) in the trio. While the alignments strongly support an 8bp homozygous deletion for both parents, the alignments for NA12878 support both a 4bp and 8bp deletion. Only 25% of the alignments for NA12878 are displayed to facilitate visualization.

## Supplementary Tables

**Supplementary Table 1:** Optimal settings for STR genotyping

| Method | Version | Explored Options | Optimal Options |
| --- | --- | --- | --- |
| freebayes | 1.0.2-6 (g3ce827d) | genotype-qualities<br>no-partial-observations<br>report-genotype-likelihood-max | genotype-qualities<br>no-partial-observations |
| GATK-HC | 3.5-0 (g36282e) | max_alternate_alleles<br>pcr_indel_model<br>kmerSize | max_alternate_alleles 25 |
| HipSTR | 0.2 | None | None |
| lobSTR | 4.0.0 | filter-mapq0<br>filter-clipped<br>max-repeats-in-ends<br>min-read-end-match | None |
| Platypus | 0.8.1 | maxVariants<br>minVarFreq | maxVariants 0.25<br>minVarFreq 0.0 |
| RepeatSeq | 0.8.2 | None | None |

**Supplementary Table 2:** Features used to build STR genotype classifiers

| **Method** | **Feature Name** | **Source<sup>1</sup>** | **Description** |
| --- | --- | --- | --- |
| freebayes | GQ | F | Genotype quality |
|  | DP | F | Genotype read depth |
|  | GLDIFF | F | Difference between genotype likelihoods of the called and next best genotypes |
|  | READRATIO<sup>2</sup> | F | Minimum ratio of read depths for called alleles |
|  | UNCALLEDFRAC<sup>2</sup> | F | Fraction of reads observed for an uncalled allele |
| GATK-HC | QD | I | Ratio of variant quality to read depth |
|  | FS | I | P-value for strand bias |
|  | ReadPosRankSum | I | Z-score for test of alt vs. ref read position bias |
|  | GQ | F | Genotype quality |
|  | DP | F | Genotype read depth |
| HipSTR | Q | F | Genotyper posterior probability |
|  | SPANDP | F | Minimum number of reads spanning either allele |
|  | DSTUTTER/DP | F | Fraction of reads with a stutter artifact |
|  | DFLANKINDEL/DP | F | Fraction of reads with an indel flanking the STR |
| lobSTR | SB | F | P-value for strand bias |
|  | Q | F | Genotype posterior probability |
|  | DP | F | Genotype read depth |
|  | DISTENDS | F | Average difference between distance of STR to read ends |
| Platypus | QD | I | Ratio of variant quality to read depth |
|  | SbPval | I | P-value for strand bias |
|  | GQ | F | Genotype quality |
|  | GOD | F | Genotype goodness of fit value |
| RepeatSeq | GQ | F | Genotype quality |
|  | DP | F | Genotype read depth |

<sup>1</sup> Describes whether the field is calculated from FORMAT fields (F) or INFO fields (I) in the variant caller's VCF
<sup>2</sup> Computed using the RO and DPR FORMAT fields

**Supplementary Table 3:** Capillary-based benchmarking of STR genotyping tools

| Method | Correct Calls<sup>1</sup> | Total Calls<sup>1</sup> | Accuracy (%) | Run Time (hrs) | Variants per STR<sup>2</sup> |
| --- | --- | --- | --- | --- | --- |
| HipSTR | 58937 | 61942 | 95.2 | 1.3 | 1 |
| GATK-HC* | 59364 | 62966 | 94.3 | 5.8 | 5.5 |
| GATK-HC | 57947 | 62966 | 92 | 6.3 | 5.5 |
| lobSTR | 54627 | 61971 | 88.2 | 0.1 | 1 |
| Platypus* | 50196 | 62937 | 79.8 | 15.7 | 4.3 |
| freebayes* | 39197 | 57349 | 63.5 | 4.8 | 2.8 |
| Platypus | 39220 | 62347 | 62.9 | 1.9 | 2.9 |
| RepeatSeq | 30289 | 52393 | 57.8 | 0.3 | 1 |
| freebayes | 9011 | 61741 | 15.7 | 3.6 | 3 |

\* Tool was run with settings optimized for accurate STR calling
1 The reported number of calls refers to the 118 samples with capillary genotypes
2 Average number of VCF records per STR region

**Supplementary Table 4:**
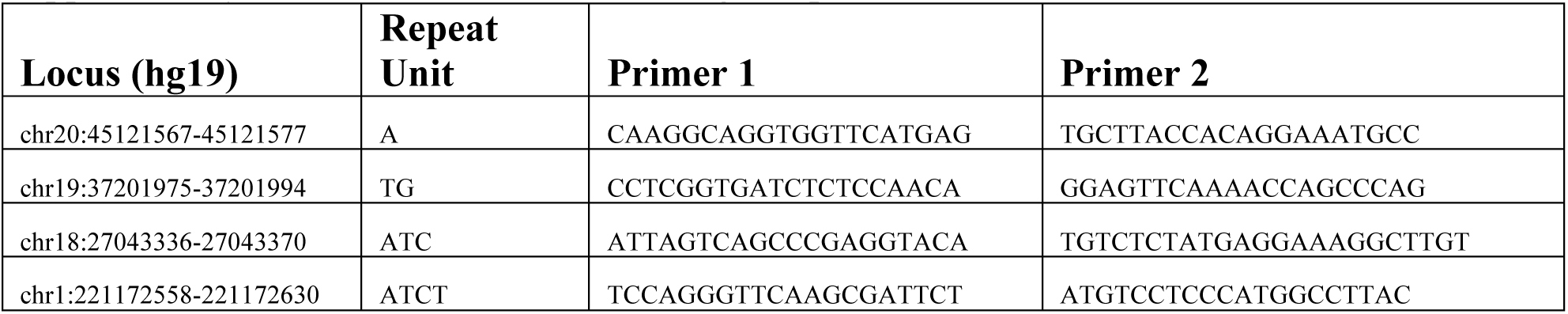
Primers for STR Sanger experiments

